# Calibration of models to data: a comparison of methods

**DOI:** 10.1101/2020.12.21.423763

**Authors:** Zenabu Suboi, Thomas J. Hladish, Wim Delva, C. Marijn Hazelbag

**Affiliations:** The South African Department of Science and Technology-National Research Foundation (DST-NRF) South African Centre for Epidemiological Modelling and Analysis (SACEMA), Stellenbosch University, Stellenbosch, South Africa; Department of Biology, University of Florida, Gainesville, FL, USA; Emerging Pathogens Institute, University of Florida, Gainesville, FL, USA; Department of Global Health, Faculty of Medicine and Health, Stellenbosch University, Stellenbosch, South Africa; International Centre for Reproductive Health, Ghent University, Ghent, Belgium; Wimmy (Pty) Ltd, Cape Town, South Africa

## Abstract

Complex models are often fitted to data using simulation-based calibration, a computationally challenging process. Several calibration methods to improve computational efficiency have been developed with no consensus on which methods perform best. We did a simulation study comparing the performance of 5 methods that differed in their Goodness-of-Fit (GOF) metrics and parameter search strategies. Posterior densities for two parameters of a simple Susceptible-Infectious-Recovered epidemic model were obtained for each calibration method under two scenarios. Scenario 1 (S1) allowed 60K model runs and provided two target statistics, whereas scenario 2 (S2) allowed 75K model runs and provided three target statistics. For both scenarios, we obtained reference posteriors against which we compare all other methods by running Rejection ABC for 5M parameter combinations, retaining the 0.1% best. We assessed performance by applying a 2D-grid to all posterior densities and quantifying the percentage overlap with the reference posterior.

We considered basic and adaptive sampling calibration methods. Of the basic calibration methods, Bayesian calibration (Bc) Sampling Importance Resampling (S1: 34.8%, S2: 39.8%) outperformed Rejection Approximate Bayesian Computation (ABC) (S1: 2.3%, S2: 1.8%). Among the adaptive sampling methods, Bc Incremental Mixture Importance Sampling (S1: 72.7%, S2: 85.5%) outperformed sequential Monte Carlo ABC (AbcSmc) (S1: 53.9%, S2: 72.9%) and Sequential ABC (S1: 21.6%, S2: 62.7%).

Basic methods led to sub-optimal calibration results. Methods using the surrogate Likelihood as a GOF outperformed methods using a distance measure. Adaptive sampling methods were more efficient compared to their basic counterparts and resulted in accurate posterior distributions. BcIMIS was the best performing method. When three rather than two target statistics were available, the difference in performance between the adaptive sampling methods was less pronounced. Although BcIMIS outperforms the other methods, limitations related to the target statistics and available computing infrastructure may warrant the choice of an alternative method.

**Author summary:** As mathematical models become more realistic, they tend to become more complex. Calibration, the process of tuning a model to better reproduce empirical data, can become dramatically more computationally intensive as model complexity increases. Researchers have responded by developing a range of more efficient, adaptive sampling calibration methods. However, the relative performance of these calibration methods remains unclear. To this end, we quantified the performance of five commonly used calibration methods. We found that adaptive sampling methods were more efficient compared to their basic counterparts and resulted in more accurate posterior distributions. We identified the best performing method, but caution that limitations related to the target statistics and available computing infrastructure may warrant the choice of one of the alternatives. Finally, we provide the code used to apply the calibration methods in our study as a primer to facilitate their application.

## Introduction

Researchers use mathematical and computer simulation models to approximate real-world processes [1–3]. Model calibration, or fitting the model to data, allows for correct estimation of uncertainty and increases the confidence that the model provides a realistic approximation to the real-world process [3, 4]. Another use of calibration is to estimate parameter values that are not available in the literature [5]. Representing parameter uncertainty allows decision-makers to assess the relative likelihood of different outcomes and not merely a point estimate of the most likely outcome. Calibrating complex models to data is often done using simulation-based calibration, which involves running the model many times [6]. This process makes simulation-based calibration challenging in terms of computational cost for models with long run-times [7].

Calibration involves running the model for different parameter combinations and comparing the produced model outputs with empirical summary statistics (targets) to identify the model parameter values that achieve a good fit to data [3, 8]. The main components of a calibration method are the targets, the parameter-search strategy, the Goodness-Of-Fit (GOF) measure, acceptance criteria, and stopping rules [4]. We distinguish Approximate Bayesian Computation (ABC) methods, using a distance measure as a GOF (e.g. Euclidean distance), from Bayesian calibration methods, using the likelihood as a GOF measure (e.g. Binomial likelihood distribution) [8, 9]. In this study, we focus on sampling algorithms as the parameter search strategy, since sampling methods obtain valid estimates of parameter uncertainty and correlations between parameters [1, 8, 10]. The basic versions of sampling algorithms, Rejection ABC and Bayesian calibration with Sampling Importance Resampling (BcSIR), draw a random sample of parameter values from the prior distribution and compute a posterior distribution [8, 11]. Researchers have proposed several adaptive sampling algorithms as more efficient parameter search strategies [3, 7, 12, 13].

Adaptive sampling algorithms improve efficiency by iteratively adapting the prior distribution such that more of the sampled parameter combinations lead to a good fit to the target statistics [7, 12, 13]. In the current study, we compare sequential Monte Carlo approximate Bayesian computation with partial least squares (AbcSmc), Adaptive population Monte Carlo ABC, hereafter called Sequential ABC - Lenormand (Seq ABC) and Bayesian calibration with Incremental Mixture Importance Sampling (BcIMIS) [7, 12, 13]. We can divide the total time consumed by calibration methods into time spent on running the model and time spent on implementing the sampling algorithm itself. Although sampling algorithms can vary substantially in run-time, the sampling algorithm’s efficiency (the number of model runs needed to achieve a certain GOF) is ultimately more important. As models become increasingly complex, the sampling algorithm run-time remains constant, while model run-time may increase arbitrarily.

A previous simulation study by Minter *et al*. found that the amount of data available for calibration affects the ability of the calibration methods to estimate the parameters [14]. The same study found that the number of retained parameter combinations, which is a function of the number of model runs performed and the tolerance of choice, influences the speed of the ABC algorithm and the accuracy of the posterior distribution. The study also found that Rejection ABC has inefficient sampling compared to the ABC Sequential Monte Carlo (ABC-SMC) algorithm [14]. Similarly, research showed that BcSIR was inefficient compared to BcIMIS [8]. Another study showed that BcIMIS outperformed both simple BcSIR and three publicly available variants of generic Markov Chain Monte Carlo (MCMC) [13]. A review of calibration methods found that the number of target statistics is often only slightly higher than the number of parameters calibrated, indicating the limited availability of target statistics to calibrate the model [15]. This review also highlighted the need for studies comparing the performance, strengths and limitations of calibration methods in scenarios differing in the number of target statistics and the number of calibrated parameters [15].

Despite this research, many questions remain around the performance of calibration methods [3, 4, 15]. Existing literature comparing the performance of calibration methods do not evaluate distance measures versus surrogate likelihoods as GOF measures and therefore do not allow us to draw general conclusions [16]. Also, previous literature does not calculate or quantify how well these adaptive sampling methods approximate the true posterior. Using a simple Susceptible-infectious-Recovered disease transmission model, we compare the performance of commonly applied basic and adaptive sampling methods, including both distance measures and surrogate likelihoods as GOF measures [15]. We assume that both the number of target statistics and the computational resources available to the researcher are limited. We highlight differences between methods concerning implementation and requirements in terms of target statistics.

## Results

### Performance for obtaining the posterior

Calibration methods differed in their ability to obtain the posterior (See Fig 1, Fig 2, and Fig 3). In Scenario 1, where the targets comprised prevalence at time 50 and prevalence at time 75, the posterior density for Rejection ABC was more dispersed than the reference posterior. The posterior densities for the other methods were closer to the reference posterior, but most methods were unable to identify *β* accurately. The posterior density for BcSIR was strongly multi-modal, in contrast to the other methods (Fig 1 and Fig A in S3 Appendix). Rejection ABC had the lowest percentage overlap of 2.3% with the reference whereas BcIMIS had the highest percentage overlap of 72.7% with the reference posterior, followed by AbcSmc with 53.9% (Fig 3). BcIMIS estimated median parameter values with credible intervals closest to their true values (Table 1). BcSIR and AbcSmc performed second best, while Rejection ABC and Seq ABC performed worst, producing biased and imprecise estimates of *β*: 0.41 (95% CI 0.17, 0.93) and 0.44 (95% CI 0.09, 0.97), respectively (Table 1). BcIMIS had a higher effective sample size (4547.48), compared to BcSIR (27.66).

**Table 1.** Median estimates of *β, γ, R*_0_, *DT* with 95% Credible Interval (CI) for both scenarios.

| | $\beta$ (95% CI) | $\gamma$ (95% CI) | $R_0$ (95% CI) | $DT$ (95% CI) |
| --- | --- | --- | --- | --- |
| True values | 0.2 | 0.02 | 10.0 | 3.85 |
| **Scenario 1 (2 targets + 60 000 model runs)** |  |  |  |  |
| Reference | 0.21 (0.17, 0.26) | 0.019 (0.016, 0.022) | 11.0 (8.3, 15.9) | 3.71 (2.86, 4.58) |
| Rejection | 0.41 (0.17, 0.93) | 0.026 (0.003, 0.077) | 19.7 (2.4, 210.8) | 1.69 (0.73, 9.88) |
| BcSIR | 0.23 (0.17, 0.39) | 0.017 (0.013, 0.021) | 13.2 (8.2, 31.0) | 3.28 (1.82, 4.50) |
| AbcSmc | 0.24 (0.17, 0.42) | 0.017 (0.013, 0.021) | 14.4 (8.1, 30.7) | 3.11 (1.73, 4.77) |
| Seq ABC | 0.44 (0.09, 0.97) | 0.014 (0.011, 0.021) | 29.7 (8.3, 78.7) | 1.77 (0.76, 4.70) |
| BcIMIS | 0.20 (0.18, 0.30) | 0.018 (0.014, 0.022) | 11.8 (7.9, 20.6) | 3.60 (2.41, 4.81) |
| **Scenario 2 (3 targets + 75 000 model runs)** |  |  |  |  |
| Reference | 0.20 (0.18, 0.23) | 0.020 (0.018, 0.023) | 9.8 (8.5, 11.4) | 3.84 (3.36, 4.37) |
| Rejection | 0.57 (0.14, 0.97) | 0.049 (0.011, 0.097) | 11.0 (6.5, 22.9) | 1.33 (0.77, 5.67) |
| BcSIR | 0.20 (0.18, 0.21) | 0.020 (0.018, 0.023) | 9.9 (7.9, 10.7) | 3.84 (3.66, 4.31) |
| AbcSmc | 0.20 (0.18, 0.22) | 0.020 (0.018, 0.022) | 10.2 (8.9, 11.8) | 3.82 (3.41, 4.30) |
| Seq ABC | 0.21 (0.17, 0.25) | 0.019 (0.017, 0.023) | 10.4 (7.8, 13.8) | 3.71 (3.02, 4.62) |
| BcIMIS | 0.20 (0.18, 0.23) | 0.020 (0.018, 0.023) | 9.9 (8.3, 11.9) | 3.84 (3.33, 4.42) |

**Fig 1.**
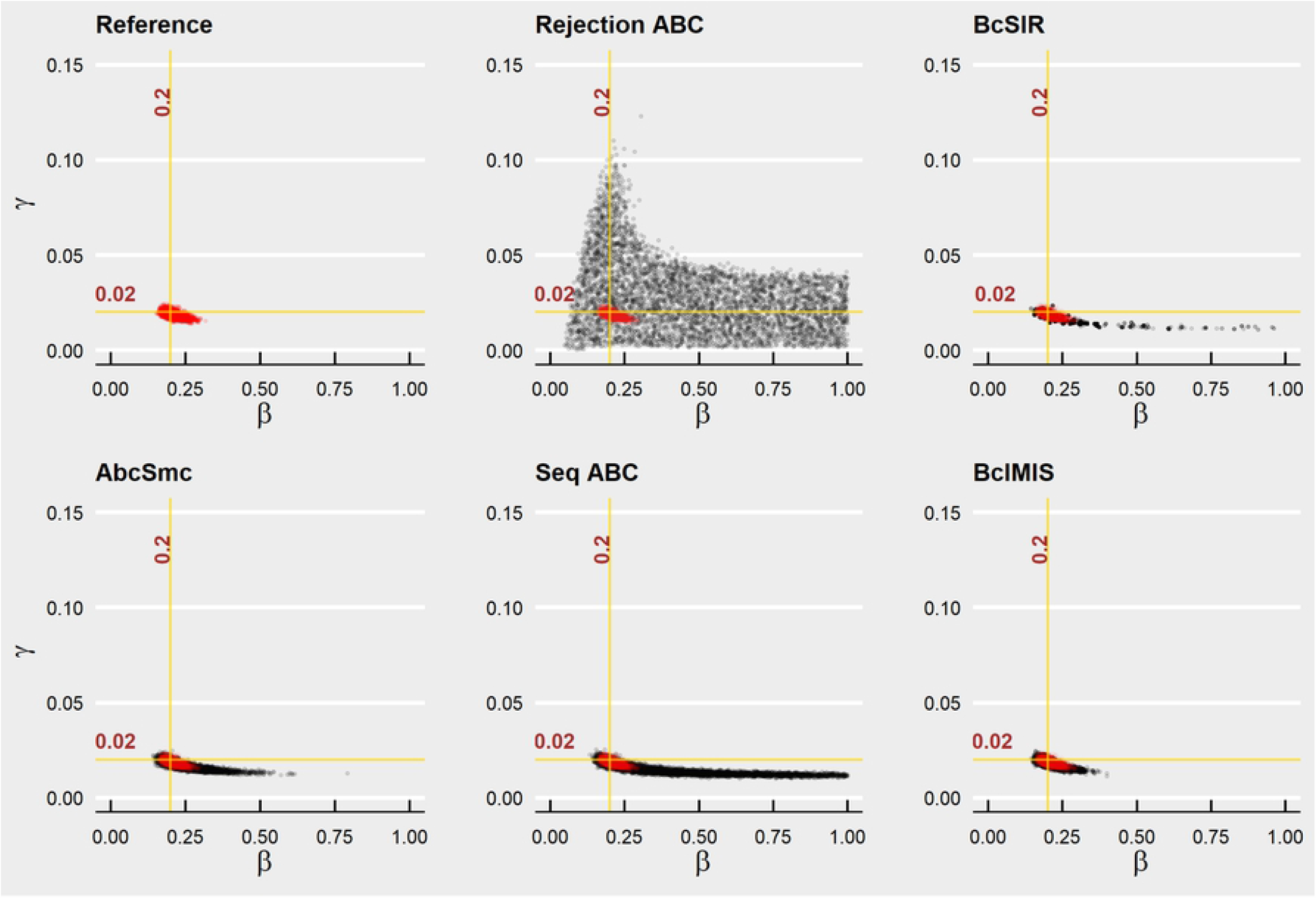
Posterior plots for Scenario 1. Plot of the posterior densities resulting from the model calibration methods (black) against the reference posterior (red) in Scenario 1. The intersection between the yellow lines represents the true parameter combination.

**Fig 2.**
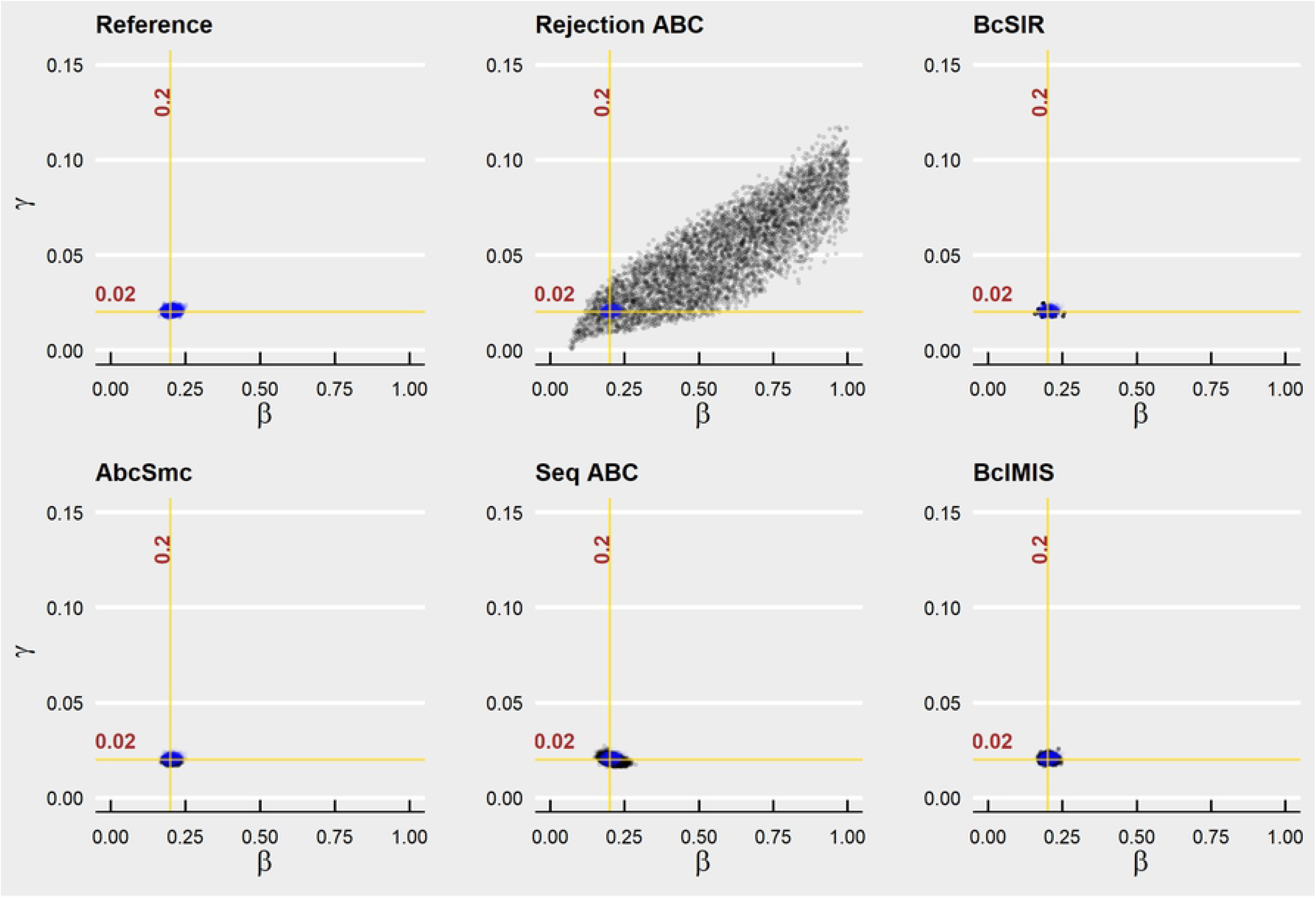
Posterior plots for Scenario 2. Plot of the posterior densities resulting from the model calibration methods (black) against the reference posterior (blue) in Scenario 1. The intersection between the yellow lines represents the true parameter combination.

**Fig 3.**
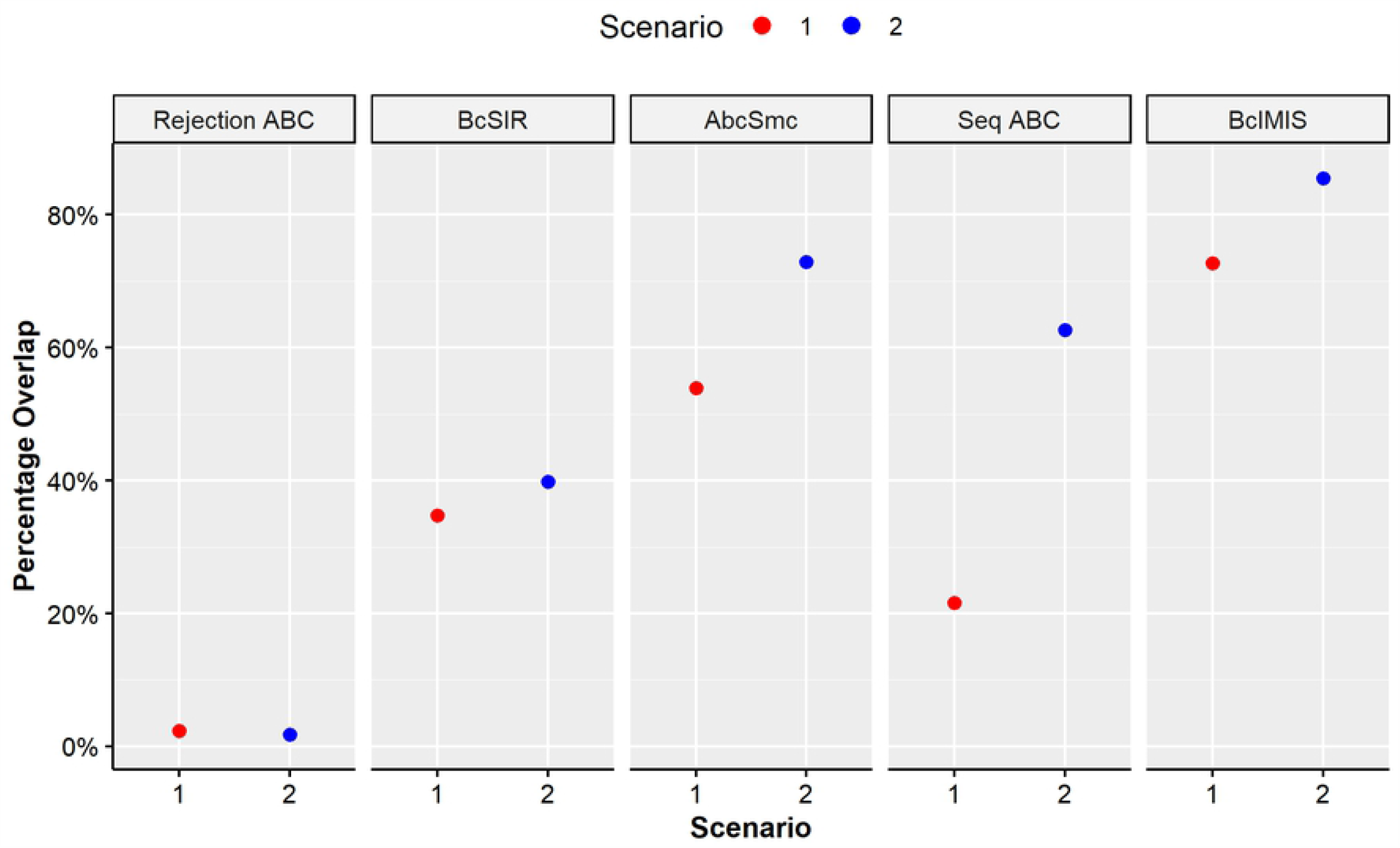
Percentage overlap plot. Plot of the percentage overlap between the posterior densities resulting from the calibration methods and the reference posterior.

In Scenario 2, where the targets consisted of prevalence at time 50, prevalence at time 75, and the peak prevalence, the posterior density for Rejection ABC was again further from the reference posterior and more spread out. In contrast, the posterior densities for the other methods were close to the reference posterior (see Fig 2 and Fig B in S3 Appendix). BcIMIS had the highest percentage overlap of 85.5% with the reference posterior, followed by AbcSmc with 72.9%. Rejection ABC had the lowest percentage overlap of 1.8% with the reference (Fig 3). All the methods, except Rejection ABC, estimated median values of *β, γ, R*_0_ and *DT* close to their true values with uncertainty similar to the reference posterior (Table 1). BcIMIS had a higher effective sample size (3552.46) compared to BcSIR (6.71).

The posterior densities for all methods in Scenario 2 were closer to the reference posterior (Fig 2 and Fig 3) than they were for Scenario 1 (Fig 1 and Fig 3), with BcIMIS being the closest for both scenarios. All methods, except Rejection ABC, estimated median values of *β, γ, R*_0_ and *DT* with less uncertainty in Scenario 2 compared to Scenario 1 (Table 1). The effective sample sizes for BcSIR and BcIMIS were higher for Scenario 1 than for Scenario 2.

## Discussion

We quantified the performance of different model calibration methods and found that adaptive sampling methods were more efficient compared to their basic counterparts and resulted in more accurate posterior distributions. Methods using the surrogate-likelihood as a GOF measure outperformed methods using a distance measure.

BcIMIS was the best performing method. When three rather than two target statistics were available, the difference in performance between the adaptive sampling methods was less pronounced. Although BcIMIS outperforms the other methods, limitations related to the target statistics and available computing infrastructure may warrant the choice of an alternative method.

BcIMIS is a Bayesian method that requires the specification and evaluation of the likelihood functions for the data underlying the target statistics. When information for target statistics is incomplete (e.g. point estimates are available but without standard errors or sample size), or when the likelihood of the observed data is intractable, the use of BcIMIS and BcSIR is no longer possible. In contrast, ABC methods only require point estimates for target statistics. When using multiple targets measured on different scales, ABC methods should employ relative distance measures to ensure equal contributions to the calibration result. An advantage of likelihood-based methods is that they allow for calculation of the effective sample size (ESS), with low values of this statistic indicating that there are only a few parameter combinations with considerable weights (e.g. as observed in Scenario 2 for BcSIR).

Complex simulation models (i.e. models with a large number of parameters and long run-times) typically require the use of high-performance computing clusters. Usability on the cluster may be another important consideration in the choice of the calibration method. The AbcSmc implementation by Hladish et al. was designed for use on a distributed cluster of compute nodes, but the same cannot be said for the implementations of Seq ABC and BcIMIS used in this analysis. If, however, the computational burden of the model calibration is small enough that a single multi-core personal computer or compute node is sufficient, Rejection ABC and seq ABC provide the convenience of a n cluster argument, allowing the user to launch model simulations in parallel on n cluster cores of the computer.

Using adaptive sampling methods potentially requires the user to restrict parameter sampling during the subsequent algorithm steps to the initial ranges of the prior distribution (e.g. disallowing negative parameter values). Seq ABC has a convenient option to do this (i.e. inside prior = TRUE). Within other methods, the use of transformations can ensure that proposed parameter values remain within the required range, e.g. sampling can be done on the logit scale, using the inverse logit to make sure that sampled parameter values stay within the 0 – 1 range. Seq ABC and BcIMIS use convergence criteria to determine when the algorithm stops. Seq ABC’s automatic convergence criterion does not allow the user to infer how many steps and how many model runs will be required before the algorithm reaches convergence. AbcSmc allows the user to specify the number of algorithm steps and the number of model runs within each step. In summary, the choice for one adaptive sampling method over another is context-dependent, conditional on computational cost the model and the availability of programming skills in the modelling group.

Our findings are consistent with some of those in previous work. Minter *et al*. similarly found that Rejection ABC was less efficient than other methods and that increasing the number of informative target statistics improves the performance of the calibration methods [14]. Our study also confirmed previous findings that BcSIR was less efficient than BcIMIS [8, 13].

A limitation of our study is the uncertainty regarding how well our results generalize to realistic calibration settings. Aspects of our experimental design might have implications for our qualitative conclusions. We used a low number of simulation runs for each of the methods (e.g. 60K in Scenario 1), while providing broad prior parameter ranges, resembling the situation in which previous research provided little information for these parameters. Using broad prior distributions may have advantaged adaptive sampling methods compared to basic calibration methods. However, we note that calibrating two parameters in scenario 1 with 60 000 simulation runs, leads to an approximate parameter sampling density of 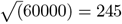, or 245*x*245 parameter combinations on the *β* by *γ* grid. As the dimensionality of parameter space increases with additional parameters, to obtain a similar parameter sampling density for calibrating three parameters would require over 14 million (245^3^) simulation runs, and for four parameters, over 3.6 billion runs. Even when prior parameter ranges are closer to the posterior density, calibrating a large number of parameters can dramatically increase the fraction of parameter space producing bad model outputs. Therefore we think that the observed increased efficiency of adaptive sampling methods over basic methods may still be an underestimation relative to realistic use cases involving more complex and imperfectly-specified models. Our use of relatively few target statistics (i.e. two to three) compared to the number of parameters we are calibrating (two) may have implications for our results. However, this design choice is in line with a recent systematic review of individual-based models that found that the number of target statistics is often limited compared to the number of parameters calibrated [15]. The last limitation is that we used default settings (e.g. convergence criterion) for most of the methods and did not explore the use of alternative method settings.

To determine the generalizability of our findings, we recommend future simulation research to explore complex models with many parameters and target statistics, calibrating more parameters to more and different types of target statistics. The performance measures used in the current study scale to these higher-dimension settings. We expect the adaptive sampling methods to outperform the basic methods exponentially as parameter space increases. Future research on adaptive sampling methods should explore the importance of sufficient exploration of the parameter space in the first set (e.g. of sequential Monte Carlo algorithms), and the optimal distribution of model runs in consecutive sets. Also, further research is required to find out whether some of the adaptive sampling methods encounter problems with the procedures to run the algorithm for more complex models.

## Methods

### Calibration Methods

We compared the following calibration methods: Rejection Approximate Bayesian Computation (Rejection ABC) [6, 9, 11], Bayesian calibration with Sampling Importance Resampling (BcSIR) [8], sequential Monte Carlo approximate Bayesian computation with partial least squares (AbcSmc) [7], Sequential Approximate Bayesian Computation - Lenormand (Seq ABC) [12], and Incremental Mixture Importance Sampling (BcIMIS) [13]. We include a description of these methods in S1 Table (see S1 Table). We provide detailed algorithms for the methods in S1 Appendix (see S1 Appendix). Table 2 provides an overview of relevant differences between the calibration methods, including parameter search strategy, GOF, parallelizability, target statistics requirements, and stopping criteria (e.g. user-specified or convergence detection).

**Table 2.**
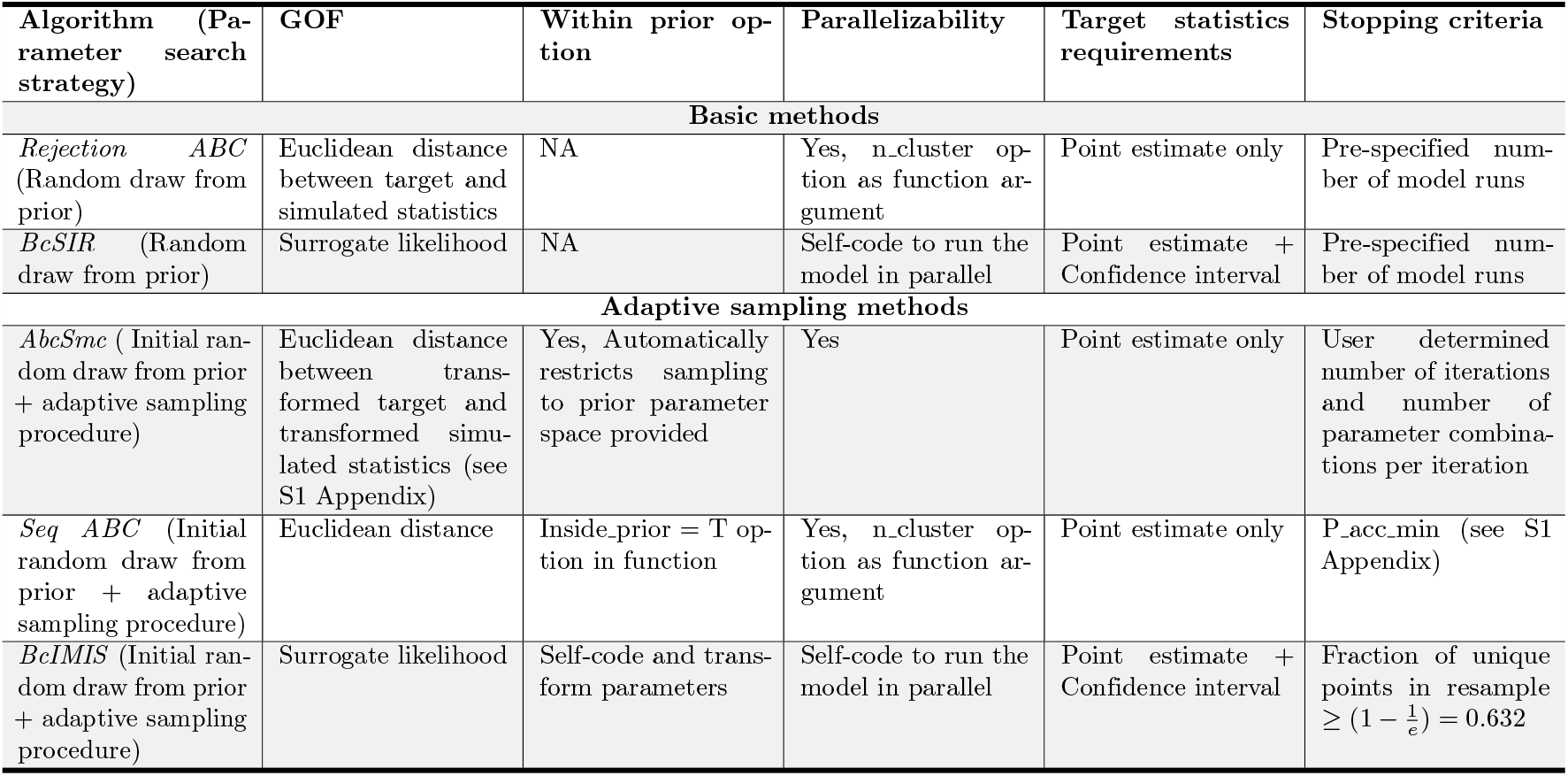
Summary of calibration methods.

### Simulation Setup

We simulated an epidemic with a known parameter combination and observed how well different calibration methods approximated a reference posterior distribution. All simulations and analyses were conducted using R version 3.6.2 (2019-12-12) [17]. The R code used to generate the simulations is available in S4 Appendix. The EasyABC package in R was used to perform simulations for Rejection ABC, and Seq ABC whereas the IMIS package was used for BcIMIS [18, 19]

### Simulation Model

We performed simulations using a simple Susceptible-Infectious-Recovered model [14]. For the model implementation we used the “SIR” function in the SimInf package in R. The SimInf package in R uses the Gillespie stochastic simulation algorithm to provide a stochastic framework for the simulation model to be implemented. The framework integrates the dynamics of infection as continuous-time Markov chains, incorporates data available, and simulates epidemics [20, 21]. We assumed correct model specification, meaning that the model used to generate the target statistics was also used to re-estimate the parameters. In the model, individuals belong to one of three compartments (i.e. health states) – Susceptible (*S*), Infectious (*I*), and Recovered (*R*). In the limit of infinite population size, the model equates to a system of three non-linear ordinary differential equations (ODEs) that relate the three compartments [22].

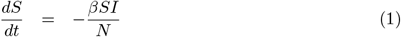

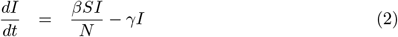

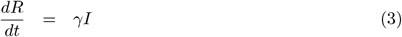

where *β >* 0 and *γ >* 0 are the rates at which individuals move from one compartment to another and *N* (*t*) = *S*(*t*) + *I*(*t*) + *R*(*t*) is the total population size which remains constant over time [23].

### Simulation Scenarios

We considered two scenarios with different numbers of prevalence-based target statistics. We define prevalence as the fraction of the population that is infectious at a given point in time. Scenario 1 (S1) considered two target statistics (prevalence at time 50 and 75) whereas scenario 2 (S2) looked at three target statistics; the two target statistics from scenario 1 plus the peak prevalence. Table 3 gives a summary of the simulation setup.

**Table 3.** Simulation Setup. Prev_*t*_ is prevalence at time *t* (or at the peak). *Methods* refers to the calibration methods we evaluated, and *reference* is the calibration we performed using a much larger Rejection ABC calibration to approximate the true posterior.

|  | Scenario 1 | Scenario 2 |
| --- | --- | --- |
| **Targets Used** | $\text{Prev}_{50} = 0.644$<br>$\text{Prev}_{75} = 0.404$ | $\text{Prev}_{50} = 0.622$<br>$\text{Prev}_{75} = 0.371$<br>$\text{Prev}_{\text{peak}} = 0.677$ |
| **Parameters used to generate targets** | $\beta = 0.2$<br>$\gamma = 0.02$ | $\beta = 0.2$<br>$\gamma = 0.02$ |
| **Initial compartmental values** | $S = 990$<br>$I = 10$<br>$R = 0$ | $S = 990$<br>$I = 10$<br>$R = 0$ |
| $\beta$ prior for methods | Uniform(0, 1) | Uniform(0, 1) |
| $\gamma$ prior for methods | Uniform(0, 0.5) | Uniform(0, 0.5) |
| **Model runs for methods** | 60K | 75K |
| $\beta$ prior for reference | Uniform(0.1, 0.4) | Uniform(0.1, 0.4) |
| $\gamma$ prior for reference | Uniform(0.01, 0.03) | Uniform(0.01, 0.03) |
| **Model runs for reference** | 5M | 5M |

The model was run once for each scenario with known parameter values *β* = 0.2, *γ* = 0.02, to obtain the prevalence values for the targets. The target statistics used in each scenario were a single realization from all possible outcomes for these values of *β* and *γ* (See Fig A and Fig B in S2 Appendix). We allowed a total of 60K and 75K model runs for scenario 1 and 2, respectively, and retained 5K parameter combinations in each. For Seq ABC, we used a stopping criterion of 0.4 as used in the Seq ABC example in the EasyABC package in R [18].

### Reference Posterior

In the limit of an infinitesimally small acceptance tolerance, Rejection ABC can be used to approximate the true posterior [9]. To increase the efficiency of obtaining the reference posterior, We reduced the width of the prior distributions for *β* and *γ* compared to the distributions used for the methods. We ran 5M simulations and accepted the 0.1% best model runs, resulting in 5K parameter combinations that form our reference posterior. To facilitate comparison with this reference posterior and to allow reliable calculation of the performance measures, all calibration methods also retained 5K parameter combinations.

### Performance Measures

#### Percentage Overlap

To compare the posterior densities of the calibration methods to the reference posterior density, we adapted the *L*_2_ measure from Lenormand et al. [12]. To this end, we created a raster to compute percentage overlaps [24]. A raster comprises a matrix of cells arranged into rows and columns to form a grid [25]. Each cell contains a number of parameter combinations. The raster was created by considering the minimum and maximum values of *β* and *γ* retained by the calibration methods. The resulting parameter space was divided into 100×100 equally sized bins with *β* values on the x-axis and *γ* values on the y-axis, leading to a grid of 100^2^ = 10000 cells. We applied the grid to each posterior density to quantify the density in each cell.

We computed the percentage overlap for each method, by summing the within cell density differences between the calibration method and the reference and subtracting from 1. Equation (4) computes the percentage overlap for each method within each scenario.

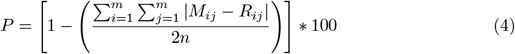

where *P* is the percentage overlap of a calibration method with the reference posterior. Summing the absolute difference between cells over the *m* = 100 rows and *m* = 100 columns of the grid approximates the overlap between the calibrated posterior and the reference posterior. In both scenarios, *M*_*ij*_ represents the cell density of row *i* and column *j* of a method in the grid, *R*_*ij*_ represents the cell density of row *i* and column *j* of the reference, *n* = 5000 is the number of parameter combinations retained by each calibration method (posterior size).

#### Quantile-Quantile Plots

We plotted the quantiles of *β* and *γ* obtained by each method against the quantiles of *β* and *γ* of the reference. Quantile-quantile plots allow us to compare distributional features, such as the presence of outliers and shifts in location.

#### Basic Reproduction Number (*R*_0_) and Epidemic Doubling time (*DT*)

We computed two derivatives of the estimated parameters; the basic reproduction number (*R*_0_), the expected number of secondary infections caused by an infectious individual in a fully susceptible population; and the doubling time (*DT*), the time it takes for the number of infectious individuals to double early in an epidemic. We computed these quantities as follows:

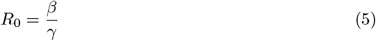

and

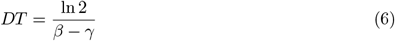

with *β - γ* being the growth rate of the epidemic.

#### Effective Sample Size

We can only calculate the effective sample size (ESS) for surrogate likelihood-based methods (i.e. BcSIR and BcIMIS). The ESS gives an estimate of the effective number of parameter combinations that ended up in our posterior, given the number of parameter combinations we sampled in total. This metric is calculated as the squared sum of the sampling weights divided by the summed squares of these weights [8]

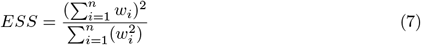

where *n* is the resample or posterior size and *w* is the sampling weight.

## Supporting information

**S1 Table. Description of calibration methods**.

**S1 Appendix. Algorithms of calibration methods**.

**S2 Appendix. Distribution of potential target statistics for both scenarios**. The targets used in our calibrations are indicated by vertical arrows.

**S3 Appendix. Close-up 2D density plots of the posteriors**. The red vertical and horizontal lines represent the true parameter values for *β* and *γ* respectively.

**S1 Fig. Quantile-Quantile plots for calibration method posteriors against the reference posteriors**. The calibration method posteriors are depicted in red (scenario 1) and blue (scenario 2) are plotted against the reference posterior (black). Coloured lines (red; scenario 1, blue; scenario 2) close to the black diagonal line indicate that the posterior distribution for the calibration method closely resembles the reference posterior.

**S4 Appendix. Simulation code**.

